# Amphetamine increases timing variability by degrading prefrontal ramping activity

**DOI:** 10.1101/2024.09.26.615252

**Authors:** Matthew A. Weber, Kartik Sivakumar, Braedon Q. Kirkpatrick, Hannah R. Stutt, Ervina E. Tabakovic, Alexandra S Bova, Young-cho Kim, Nandakumar S. Narayanan

## Abstract

**Background:** Amphetamine is a commonly abused psychostimulant that increases synaptic catecholamine levels and impairs executive functions. However, it is unknown how acute amphetamine affects brain areas involved in executive control, such as the prefrontal cortex. We studied this problem in mice using interval timing, which requires participants to estimate an interval of several seconds with a motor response. Rodent prefrontal cortex ensembles are required for interval timing. We tested the hypothesis that amphetamine disrupts interval timing by degrading prefrontal cortex temporal encoding.

**Methods:** We first quantified the effects of amphetamine on interval timing performance by conducting a meta-analysis of 11 prior rodent studies. We also implanted multielectrode recording arrays in the dorsomedial prefrontal cortex of 7 mice and then examined the effects of 1.5 mg/kg *_D-_*amphetamine injected intraperitoneally on interval timing behavior and prefrontal neuronal ensemble activity.

**Results:** A meta-analysis of previous literature revealed that amphetamine produces a large effect size on interval timing variability across studies but only a medium effect size on central tendencies of interval timing. We found a similar effect on interval timing variability in our task, which was accompanied by greater trial-to-trial variability in prefrontal ramping, attenuated interactions between pairs of ramping neurons, and dampened low-frequency oscillations.

**Conclusions:** These findings suggest that amphetamine alters prefrontal temporal processing by increasing the variability of prefrontal ramping. Our work provides insight into how amphetamine affects timing-related brain activity, which may be useful in developing new neurophysiological markers for amphetamine use and novel treatments targeting the prefrontal cortex.

## Introduction

Amphetamine has complex effects on the central nervous system, including disruptions to vesicular storage, metabolism, and reuptake, which broadly increase release and synaptic levels of catecholamines such as dopamine and norepinephrine (1–3). While these mechanisms can potentially have myriad effects, it is unclear how amphetamine affects neurons and neuronal networks *in vivo*. This is particularly salient because amphetamine is a clinically prescribed psychostimulant that is commonly abused and can degrade executive functions in humans (4–8). Understanding the neuronal and network-level effects of amphetamine is important, as this knowledge could lead to neurophysiological markers of amphetamine use as well as new interventions that minimize abuse and maximize therapeutic potential.

We studied this problem by investigating how amphetamine affects neuronal and network activity during interval timing. This cognitive paradigm requires participants to estimate an interval of several seconds by making a motor response and involves executive functions, such as working memory for temporal rules and attention to the passage of time. Interval timing is ideal to investigate neuronal and network effects of amphetamine because it can be studied in both humans and rodent models (9), and it requires highly conserved forebrain circuits in the prefrontal cortex (10,11). Neurotransmitters that are affected by amphetamine, such as dopamine, reliably affect interval timing across species (12–14).

Past work by our group and others has established that prefrontal cortex neuronal ensembles encode temporal information (13–21) through time-dependent linear changes in firing rates over an interval. Importantly, this pattern of ramping activity is dependent on intact prefrontal cortex dopamine (13,14,16,22). These data led to the hypothesis that amphetamine disrupts interval timing by degrading temporal encoding by prefrontal neurons. We tested this idea by first performing a meta-analysis of past studies that investigated the effect of amphetamine on interval timing in rodents. We then examined the effects of amphetamine on interval timing behavior and prefrontal neuronal ensemble activity by recording prefrontal neuronal ensembles during interval timing in mice. We found evidence of increased interval timing variability and increased neuronal variability among prefrontal ensembles that degrades temporal processing. These data capture how amphetamine modulates executive processing in the prefrontal cortex.

## Methods

We reviewed previous literature to assess the effects of acute amphetamine administration on interval timing in rodents. To identify the acute effects of amphetamine on interval timing behavior, we conducted an electronic search of PubMed and PsychInfo using the terms “amphetamine” and “interval timing,” which yielded 69 peer-reviewed manuscripts; one additional manuscript was found while reviewing relevant manuscripts for a total of 70 manuscripts. Two authors independently screened all manuscripts (MAW and BQK) to identify peer-reviewed original research investigating the effect of acute amphetamine on interval timing performance in rodents. This screening process resulted in 15 peer-reviewed manuscripts. Features collected from each study were: 1) article title; 2) authors; 3) publication year; 4) amphetamine dose and route of administration; 5) time between amphetamine administration and interval timing behavior; and 6) type of interval timing behavior. We were interested in two key dependent variables of interval timing behavior: 1) *timing precision*: the measure of timing variability, or the spread of timed responses (measured using Weber fraction; peak spread, etc.); and 2) *timing accuracy*: the measure of a left or right shift that indicates early or late timing, respectively. Timing accuracy represents the central tendency of timed responses (measured using peak response time, bisection point, etc.). We extracted these variables along with the standard deviation and/or standard error of the mean (SEM) and the number of subjects. Four manuscripts (23–26), were removed because either the mean, standard deviation, or SEM could not be extracted, leaving a total of 11 manuscripts for analysis.

For each of the 11 studies, we calculated the Cohen’s *d* effect size for measures of timing precision and timing accuracy (27). In meta-analyses, the Cohen’s *d* effect size is a unitless measure that facilitates comparisons across studies with diverse methods. Of these 11 manuscripts, several studies reported the effects of multiple doses of amphetamine (7/11 studies), some reported the effects of amphetamine on different types of interval timing tasks (3/11 studies), and one reported the effects of amphetamine on different rodent strains (1/11). Therefore, we calculated multiple effect sizes, when available, and included these in the meta-analysis (28). Effect sizes were adjusted so that decreased timing variability or a leftward shift in timing were reflected by negative values, and increased variability or a rightward shift in timing were reflected by positive values. We excluded individual Cohen’s *d* values greater than or less than ±4 (n = 1 of 2 data points from Fowler & Pinkston, 2009 (29)), as in prior work as such values would have an outsize effect (28). Finally, we used a fixed-effects meta-regression model to determine whether acute amphetamine produced a larger effect on either interval timing variability or interval timing accuracy. Statistical analyses were completed using R statistical software (v 4.3.2; R Core Team) and the *metafor* meta-analysis package for R (30,31).

### Mice

All procedures were approved by the Institutional Animal Care and Use Committee (IACUC) at the University of Iowa. All methods were performed in accordance with the relevant guidelines and regulations (Protocol #3052039). We included a total of 7 mice (3 C57BL/6J (3 females) and 4 DAT^IRES^-Cre (2 females, 2 males)). C57BL/6J mice were obtained from Jackson Labs at 12–14 weeks of age and acclimated to our animal holding facility for at least 1 week prior to starting experiments. DAT^IRES^-Cre mice (Jackson Labs #006660, Bar Harbor, ME) were bred in-house and maintained on a C57BL/6J background. All mice were singly housed on a 12-hour light/dark cycle with *ad lib* access to laboratory rodent chow and water, until starting food restriction to ∼85% of free-fed weight. Interval timing training described in detail below began approximately 1 week after starting food restriction.

### Stereotaxic surgery

Mice were anesthetized with 4.0% isoflurane at 400 mL/min and maintained on 1.5%–3.0% isoflurane at 120 mL/min (SomnoSuite, Kent Scientific, Torrington, CT, USA). Craniotomies were drilled above the frontal cortex (AP +1.8, ML +/- 0.3; counterbalanced left vs. right across mice), and 4×4 (1 mm^2^) multielectrode recording arrays (MicroProbes, Gaithersburg, MD) were implanted (DV -1.8). At least three more craniotomies were drilled to insert skull screws (to anchor the headcap assemblies) and ground electrode arrays; these were sealed with cyanoacrylate (“SloZap,” Pacer Technologies, Rancho Cucamonga, CA) and accelerated with “ZipKicker” (Pacer Technologies) and methyl methacrylate (AM Systems, Port Angeles, WA). Following surgery, mice were allowed to recover for 1 week before restarting food restriction and interval timing training.

### Mouse interval timing

We used a mouse-optimized interval timing switch task (described in detail previously (32–37)), in which mice *respond* by switching nosepoke ports after ∼6 seconds to receive a food reward. Briefly, mice are trained in standard sound-attenuated operant chambers (MedAssociates, St. Albans, VT). Sessions last 90 minutes, with ∼50% of trials consisting of a “short” 6-second interval, and the remaining trials consisting of a “long” 18-second trial. Importantly, short and long trials have identical audio and visual cues and are self-initiated by a response at the back nosepoke. In each trial type, mice must respond at a designated short or long nosepoke port after 6 or 18 seconds, respectively, to receive a 20-mg sucrose reward (BioServ, Flemington, NJ). When a short trial is initiated, mice begin by responding at the designated short trial nosepoke, and a reward is delivered for the first response after 6 seconds; these trials are not analyzed. On long trials, a reward is delivered for the first response at the designated long trial nosepoke after 18 seconds. Since cues are identical for both short and long trials, the optimal strategy is for mice to start responding at the designated short trial nosepoke, but if enough time passes with no reward delivered, the mice switch to the designated long trial nosepoke (Fig. 2A). The *switch response time* is defined as the moment when the mouse leaves the short trial nosepoke and is a measure of the rodent’s internal estimate of time, as in other interval timing tasks (38,39). Only the long trials in which mice make an appropriate switch are analyzed. After achieving optimal performance in the interval timing task, which takes approximately 4 weeks, mice are acclimated to handling and the electrophysiological recording procedures described in detail below.

### Amphetamine administration

We examined the effect of acute amphetamine on interval timing behavior and frontal cortex neuronal ensembles. On Day 1, all 7 mice were injected intraperitoneally (IP) with 0.9% saline, 20 minutes before interval timing sessions and recordings were performed. On Day 2, the same mice were injected with *_D-_*amphetamine (1.5 mg/kg dissolved in sterile 0.9% saline, IP; Millipore-Sigma A5880, Burlington, MA), 20 minutes before interval timing sessions and recordings were performed.

### Neurophysiological recordings and neuronal analyses

Recordings of frontal cortex neuronal ensembles were collected through a multielectrode recording system (Open Ephys, Atlanta, GA) during interval timing sessions after saline (Day 1) and amphetamine (Day 2) administration. Offline spike sorting software, Offline Sorter™ (Plexon, Dallas, TX), was used to remove artifacts and classify single units using principal component analysis (PCA) and waveform shape. Importantly, sessions that were recorded separately from mice that had been administered saline (Day 1) and amphetamine (Day 2) were sorted together in the same PCA space, allowing us to examine the effects of saline vs. amphetamine within individual neurons across sessions. Single units were defined as those with 1) a consistent waveform shape; 2) a separable cluster in PCA space; and 3) a 2-ms or longer refractory period in the inter-spike interval histogram. Spike activity was calculated using kernel density estimates of firing rates across the interval (-4 seconds before trial start to 22 seconds after trial start), binned at 0.25 seconds with a bandwidth of 1.

As in our previous work, we defined time-related ramping activity as a monotonic change in firing rate over the whole interval (13,15,34,39). To estimate the average slope and the confidence interval (CI) of firing rates across all trials, we fit generalized linear regression models (GLM; *fitglme* in MATLAB) for each neuron. A significant linear fit (i.e., time-related ramping) was tested using *anova* in MATLAB; only neurons with a *p* value of <0.05 after correction using Benjamini-Hochberg false discovery rate were considered to exhibit time-related ramping.

### Joint peristimulus time histograms

To investigate functional interactions between pairs of frontal cortex neurons, we calculated joint peristimulus time histograms (JPSTHs), described in detail in our previous work (15,40). Briefly, JPSTHs were computed using 1-second bins from -4 seconds before trial start to 22 seconds after trial start; bin size was chosen based on our previous work showing maximal interactions using 1-second bins (15). To account for modulations in firing rates, we shuffled trial order for each pair of neurons, creating a shift-predicted matrix. We then subtracted the shift-predicted matrix from the raw JPSTH matrix. This analysis was first done with unique pairs of ramping or non-ramping neurons in mice that were administered saline on Day 1. We then calculated JPSTHs using the exact same pairs of neurons in mice that received amphetamine administration on Day 2, regardless of whether a neuron displayed or did not display significant time-related patterns of activity.

### Time-frequency analyses

Our prior work has highlighted the importance of dopamine and D1-receptors for frontal cortical low-frequency oscillations (1–8 Hz) during interval timing (13–16,41). Similar to our prior work (13,14,16,41,42), we computed spectral measures by multiplying the fast Fourier transformed (FFT) power spectrum of single-trial data with the FFT power spectrum of a set of complex Morlet wavelets (defined as a Gaussian-windowed complex sine wave: 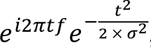, where *t* = time and *f* = frequency). These wavelets increased in 100 logarithmically-spaced steps from 1–50 Hz, and the cycles of each frequency band increased from 3–12 cycles between 1–50 Hz, taking the inverse FFT. To allow direct comparison of effects across frequency bands and treatment conditions, power was then converted to a decibel (dB) scale (10*log10[power(t)/power(baseline)]). Because the interval timing task is self-initiated, the pre-trial baseline was defined at -3 seconds to -2 seconds before trial start. As in our prior work, we used a *t*-test to find differences between treatment conditions with cluster sizes of at least 500 pixels (14,16,42–44).

### Histological procedures

Mice were anesthetized with IP injections of ketamine (100 mg/kg) and xylazine (10 mg/kg) in preparation for transcardial perfusion. Perfusions were performed using ice-cold phosphate buffer solution (PBS) and 4% paraformaldehyde (PFA). Brains were removed and fixed in 4% PFA for 24 hours, transferred to 30% sucrose for 48 hours, and then frozen in cryoprotectant at -80 ℃. A cryostat (Leica Biosystems, Deer Park, IL) was used to collect 40-µm horizontal sections of the frontal cortex. Selected slices of prefrontal cortex tissue were stained to visualize tyrosine hydroxylase fluorescence, according to methods described in detail previously (33,37). All images were captured using VS-ASW-S6 imaging software (Olympus, Center Valley, PA) to confirm histological location of multielectrodes in the prefrontal cortex.

### Statistics

All data were analyzed with custom code written in MATLAB and R, and the statistical approaches were reviewed by the Biostatistics, Epidemiology, and Research Design Core (BERD) in the Institute for Clinical and Translational Science (ICTS) at the University of Iowa; all code and data are available at http://narayanan.lab.uiowa.edu/article/datasets. The two primary measures of interval timing performance were quantified by: 1) the switch response time coefficient of variance (CV) of all trials performed during an interval timing session and 2) the mean switch response time of all trials performed during an interval timing session. CV was calculated as the standard deviation divided by mean and is expressed as a percentage. We also analyzed the average number of nosepoke responses per switch trial, average nosepoke response duration, average time from the final short nosepoke to the first long nosepoke during a switch response, and the number of switch trials. Differences between interval timing performance, neuronal ensemble recording data, and local field potentials (LFPs) were computed using a one-way repeated measures ANOVA (*anova_test* in R *rstatix* package) or using ANOVA from linear models (*lmer* in R *lmerTest* package). We accounted for animal-specific variance in all analyses, set statistical significance at alpha 0.05, and used Cohen’s *d* to calculate effect size for reliable effects.

## Results

### Meta-analysis of the effect of acute amphetamine on interval timing

We conducted a meta-analysis of results from 11 previous studies to summarize the effect of acute amphetamine on interval timing in rodents (29,32,45–53). We first ran a meta-analysis to determine the effect of acute amphetamine on interval timing variability (timing precision) and found a significant increase in mice that had been administered amphetamine compared to controls, with a Cohen’s *d* effect size of 0.95 (95% CI 0.65–1.26, *p* = 9.4 × 10^-10^; Fig. 1). We then ran a separate meta-analysis of the same studies to examine early or late shifts in interval timing (timing accuracy) and found a significant decrease in response times, i.e., a leftward shift in timed responses, among mice that had been administered amphetamine compared to controls, with a Cohen’s *d* effect size of -0.43 (95% CI -0.67 – -0.19, *p* = 0.0005; Fig. 1). Thus, these data indicate a large effect size of acute amphetamine on timing precision and a medium effect size on timing accuracy.

**Figure 1.**
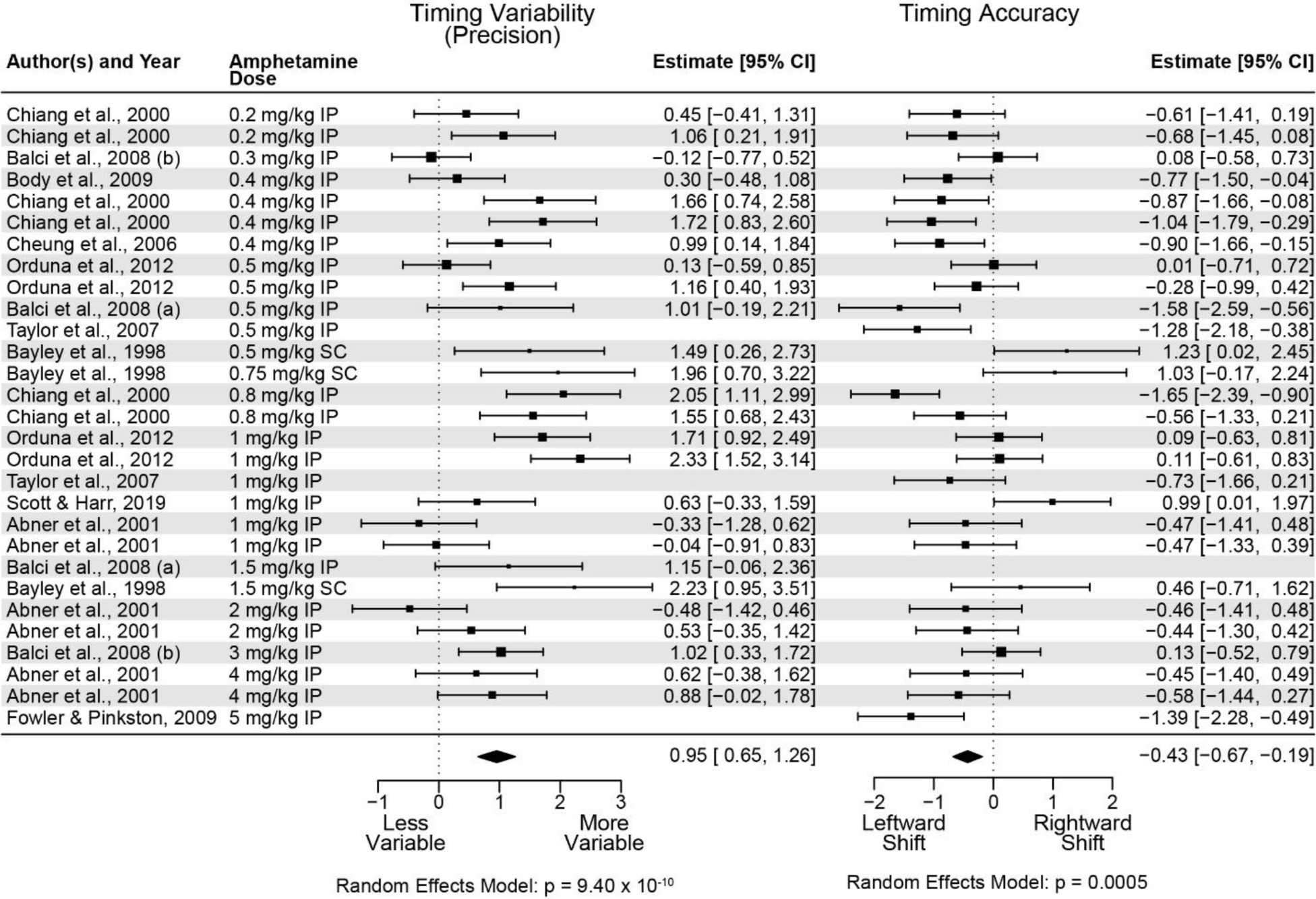
Meta-analysis of 11 prior studies investigating the effect of amphetamine on interval timing behavior in rodents. Acute amphetamine produced a significant increase in timing variability (timing precision; large effect size = 0.95; data from 26 Cohen’s *d* effect sizes in 11 rodent studies) and a significant leftward shift in timing accuracy (medium effect size = -0.43; data from 28 Cohen’s *d* effect sizes in 11 rodent studies).

To determine whether amphetamine administration produced a statistically larger effect on timing variability compared to timing accuracy, we used a fixed-effects meta-regression model. Because a leftward shift in timing accuracy is associated with a negative Cohen’s *d* value, we compared the absolute value of timing accuracy effect size to the timing variability effect size. The meta-regression model indicated that amphetamine had a significantly larger effect on timing variability compared to timing accuracy (estimate = 0.634; z = 6.527; 95% CI 0.44 – 0.82, *p* = 6.69 × 10^-11^).

### Amphetamine alters interval timing behavior

Next, we examined the effect of acute amphetamine on interval timing behavior in an interval timing task in which mice were rewarded for switching from one nosepoke port to another after estimating that enough time had passed at the first nosepoke port without reward (32). This interval timing task is described in detail in our past work (33–36). We define the *switch response time* as the moment when a mouse exits the short nosepoke prior to responding at the long nosepoke, as this response is guided by the temporal control of action; on Day 1, all mice were injected with 0.9% saline, and on Day 2, the same mice were injected with *_D-_*amphetamine (Fig. 2A). Response density for short responses, the switch response, and long responses are shown in Fig. 2B, and the cumulative distribution of a switch for each mouse is shown in Fig. 2C. We found that acute amphetamine reliably increased the switch response time coefficient of variance (CV; measure of timing precision; saline: 29.48 ± 2.55% vs. amphetamine: 34.74 ± 2.57%; F(_1,6_) = 26.18, *p* = 0.002; Cohen’s *d* = 0.775; Fig. 2D), and trended to decrease the mean switch response time (measure of timing accuracy; saline: 10.21 ± 0.32 (mean ± SEM) seconds vs. amphetamine: 8.45 ± 0.48 seconds; F(_1,6_) = 5.79, *p* = 0.053; Cohen’s *d* = 1.657; Fig. 2E) Amphetamine did not change the mean number of responses per trial (saline: 8.36 ± 1.89 responses vs. amphetamine: 7.15 ± 2.25 responses; F(_1,6_) = 1.75, *p* = 0.234; Fig. S1A) but did change the short response duration (saline 0.32 ± 0.03 seconds vs. amphetamine 0.27 ± 0.02 seconds; F(_1,6_) = 10.71, *p* = 0.017; Cohen’s *d* = 0.769; Fig. S1B) and long response duration (saline 0.34 ± 0.03 seconds vs. amphetamine 0.29 ± 0.02 seconds; F(_1,6_) = 6.15, *p* = 0.048; Cohen’s *d* = 0.824; Fig. S1C). Further, amphetamine trended to change the time from the final short nosepoke to the first long nosepoke during a switch response (saline: 5.29 ± 0.81 seconds vs. amphetamine: 6.72 ± 0.88 seconds; F(_1,6_) = 4.97, *p* = 0.067; Fig. S1D) but did not reliably change the number of switch trials (saline: 19.86 ± 2.64 trials vs. amphetamine: 17.00 ± 2.01 trials; F(_1,6_) = 2.79, *p* = 0.146; Fig. S1E).

**Figure 2.**
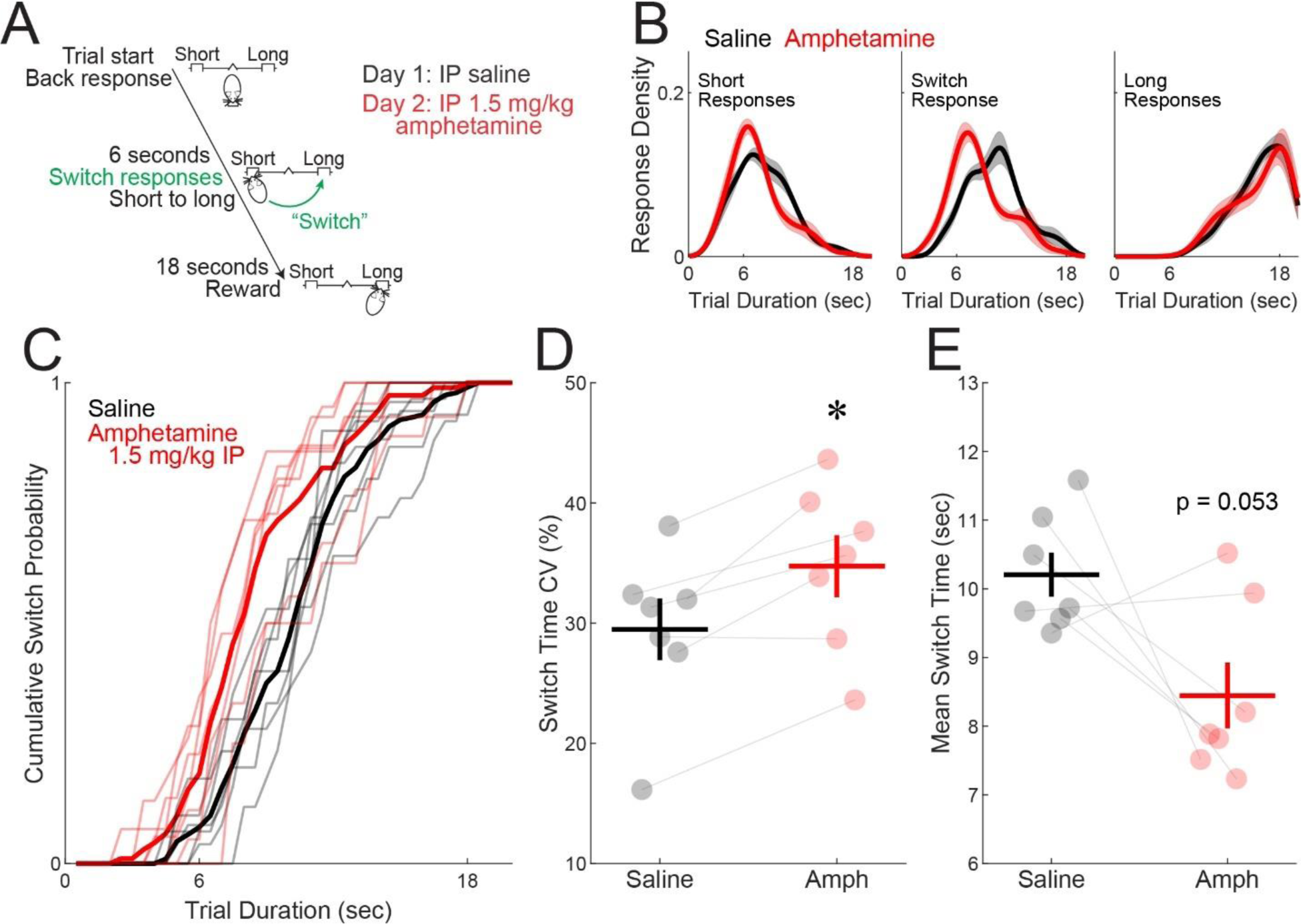
Amphetamine alters interval timing behavior. **(A)** Mice were trained on an interval timing task in which they must switch nosepoke ports based on an internal estimation of how much time has passed. In well-trained mice, we recorded from frontal cortex neuronal ensembles following intraperitoneal (IP) injection of saline (Day 1) and IP injection of 1.5 mg/kg amphetamine (Day 2). **(B)** Response density for short responses (left), switch response (middle), and long responses (right) under saline (black) and amphetamine (red). Shaded area is standard error. **(C)** Cumulative switch probability for each mouse and average cumulative switch probability. **(D)** Switch response time coefficient of variance (CV; interval timing precision) and **(E)** mean switch response time (interval timing accuracy). Each dot represents a single mouse; 5 females (2 DAT^IRES^-Cre, 3 C57BL/6J) and 2 males (2 DAT^IRES^-Cre). The horizontal line represents the mean, and the vertical line represents standard error. * *p* < 0.05

### Amphetamine alters time-related ramping patterns of neuronal activity

We recorded neuronal ensembles from the rodent dorsomedial prefrontal cortex (Fig. 3A), according to methods described in detail previously (13,14,16,39,41,54). We sorted and matched 99 neurons across saline and amphetamine interval timing sessions. Neuron waveform shape across all sessions was highly correlated (R^2^ = 0.996 ± 0.0016; Fig. 3B), and inter-spike intervals did not significantly differ (saline: 0.32 ± 0.04 seconds vs. amphetamine: 0.34 ± 0.05 seconds; F(_1,98_) = 0.45, *p* = 0.502). We removed 9 neurons with firing rates of <0.5 Hz, leaving 90 matched neurons for analysis.

**Figure 3.**
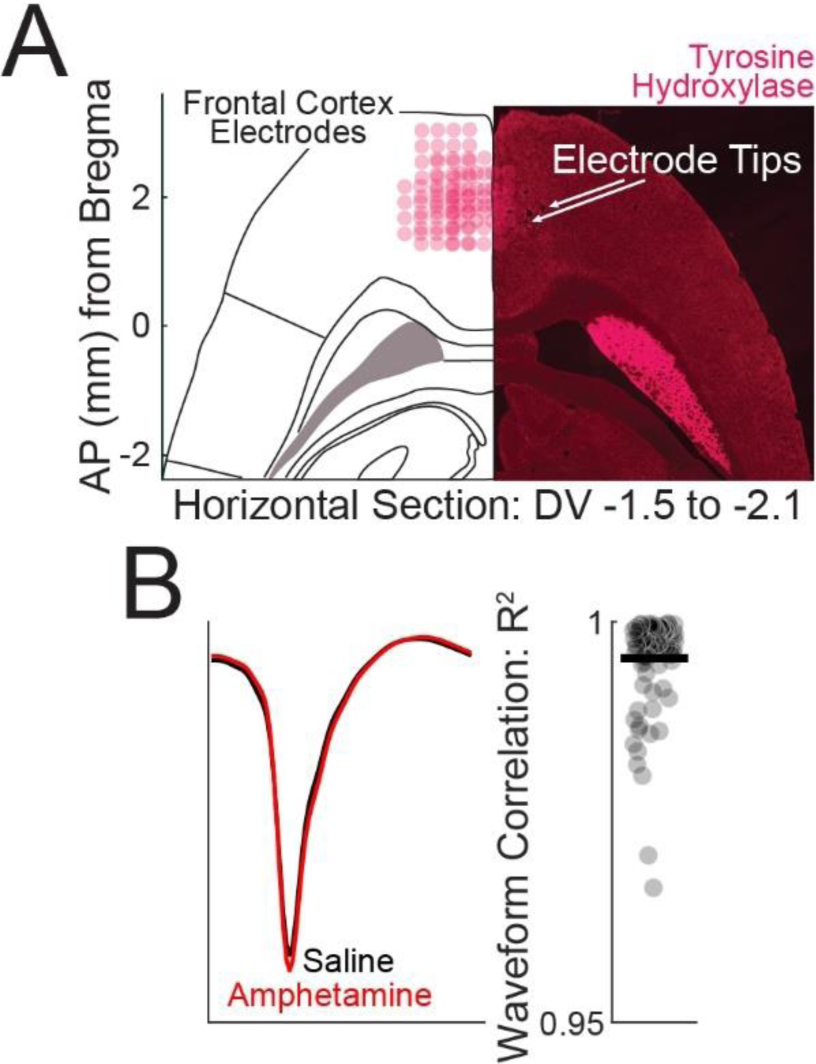
Single-unit electrophysiology recordings in the frontal cortex. **(A)** Horizontal section depicting electrode reconstruction from 7 mice. **(B)** Left: example waveform averaged across all action potentials from one prefrontal cortex neuron after saline (black) or amphetamine (red). Right: waveforms from identified neurons were highly correlated across recording sessions. Each dot represents a single neuron. The horizontal line represents the mean. Data from 99 well-isolated and sorted neurons in 7 mice.

We constructed peri-event time histograms (PETHs) using kernel density estimates and z-score normalization of firing rates between -4 seconds and 22 seconds from the trial start for both treatment conditions (Fig. 4A & B). We then used data-driven principal component analysis (PCA) to quantify differences in patterns of neuronal activity between sessions. PCA was computed using neuronal ensembles from saline and amphetamine sessions. As in our past studies, PCA identified time-dependent ramping activity as PC1 (13,15,16,34,39,54), which explained ∼34% of the variance among prefrontal cortical neurons (Fig. 4C & D). Interestingly, we found that amphetamine administration shifted PC1 scores (saline: 0.70 ± 0.52 vs. amphetamine: -0.70 ± 0.52; F(_1,89_) = 6.14, *p* = 0.015; Cohen’s *d* = 0.283; Fig. 4E). These data show that time-related ramping activity, as measured by PC1, is changed with acute amphetamine. Although PCA is data-driven, it is calculated from neuronal averages, which minimize variance across ensembles. There could be two reasons for decreased ramping activity: 1) a decrease in ramp slope over the 18-second interval during interval timing switch trials or 2) an increase in trial-to-trial variability.

**Figure 4.**
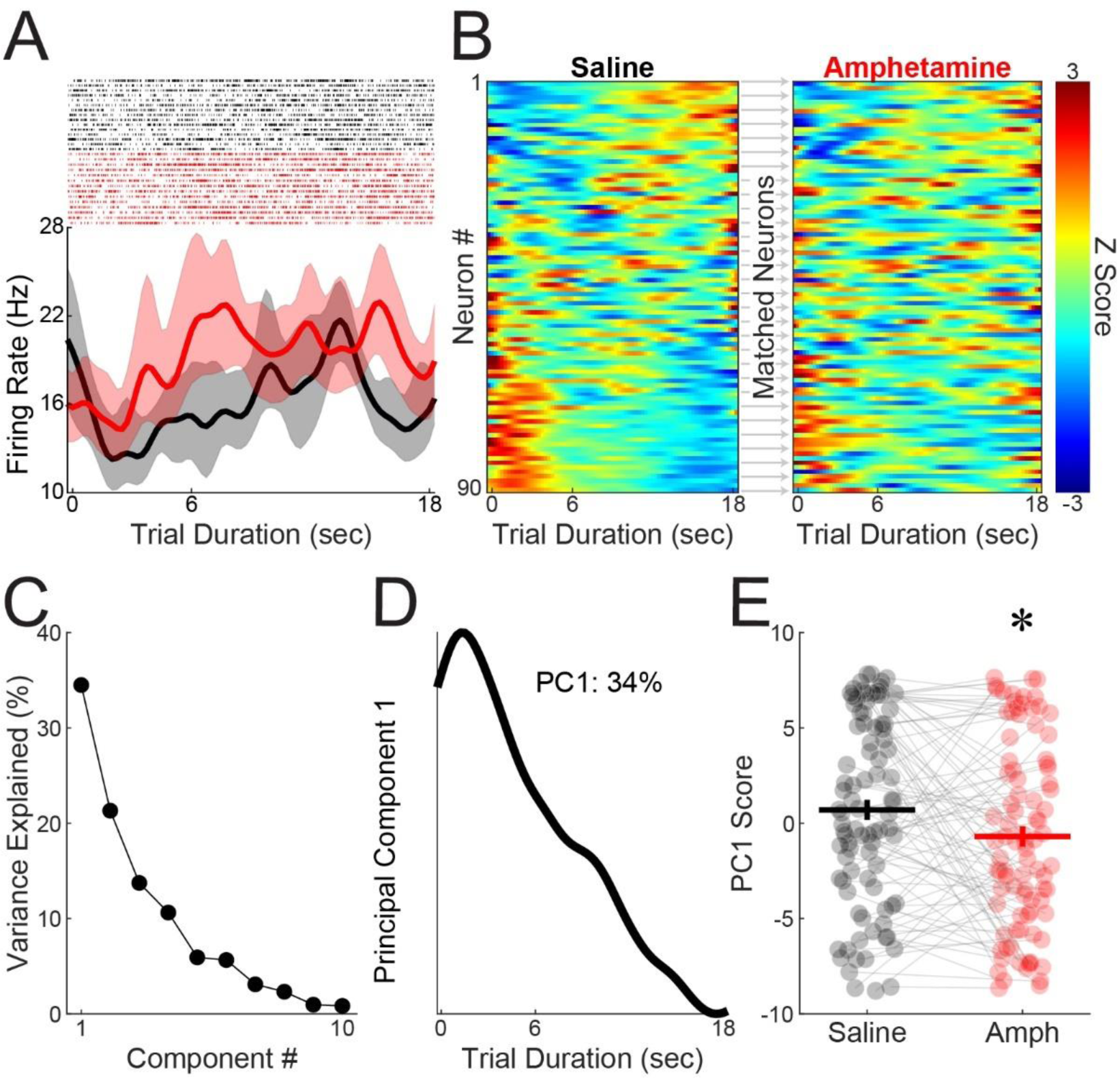
Amphetamine alters temporal processing in the frontal cortex. **(A)** Top: example peri-event raster during interval timing sessions with saline (black) and amphetamine (red). Bottom: estimated mean firing rate across all trials; shaded area is the bootstrapped 95% confidence interval. **(B)** Peri-event time histograms showing single neurons (n = 90) from 7 mice with saline (left) and amphetamine (right); neurons were matched across sessions based on waveform and inter-spike interval (see Fig. 3). Each row represents a peri-event time histogram (PETH) across all switch trials, binned at 0.25 seconds, smoothed using kernel density estimates, and z-scored. Colors indicate z-scored firing rates. **(C–D)** Principal component analysis (PCA) of frontal cortex ensembles exhibited time-dependent ramping, which explained ∼34% of population variance. **(E)** PC1 scores were significantly shifted with amphetamine administration. The horizontal line represents the mean, and the vertical line represents standard error. * *p* < 0.05. Data from 90 well-isolated and sorted neurons in 7 mice.

We tested these ideas with general linear models (GLMs) that quantify neuronal ramping activity on a trial-by-trial basis according to the equation:

#### *Firing Rate* ∼ *Time* * *Responses*

This analysis includes neuronal activity in individual trials and is not influenced by trial averaging. GLMs that were corrected for multiple comparisons identified 41 neurons with significant time-dependent ramping activity under the saline condition (∼46% of all neurons; Fig. S2A). Of these, there were 15 up-ramping neurons (∼17% of all neurons) and 26 down-ramping neurons (∼29% of all neurons), which is similar to our prior work (15). Under the amphetamine condition, we found that 37 neurons ramped (∼41% of all neurons; Fig. S2A); however, of the 41 neurons that ramped under the saline condition, only 22 of those neurons still ramped under the amphetamine condition (Fig. S2A). We then investigated how amphetamine changes ramping activity in these neurons. First, for the 41 time-related ramping neurons under the saline condition, trial-by-trial mean firing rate did not significantly change as result of amphetamine administration (saline: 11.27 ± 1.50 Hz vs. amphetamine: 12.45 ± 1.59 Hz; F(_1,40_) = 2.79, *p* = 0.10; Fig. 5A). Furthermore, the trial-by-trial slope did not significantly change with amphetamine administration (saline: -0.10 ± 0.04 spikes/second vs. amphetamine: -0.078 ± 0.04 spikes/second; F(_1,40_) = 0.51, *p* = 0.479; Fig. 5B). Strikingly, the confidence intervals (CIs) of trial-by-trial slope estimates significantly increased with amphetamine administration, suggesting that ramping slope became more variable with amphetamine (saline: 0.15 ± 0.01 vs. amphetamine: 0.17 ± 0.02; F(_1,40_) = 13.10, *p* = 0.0008; Cohen’s *d* = 0.273; Fig. 5C). In the amphetamine condition, the CI size increased in 29 of the 41 ramping neurons, with 22 neurons increasing by more than 10%.

**Figure 5.**
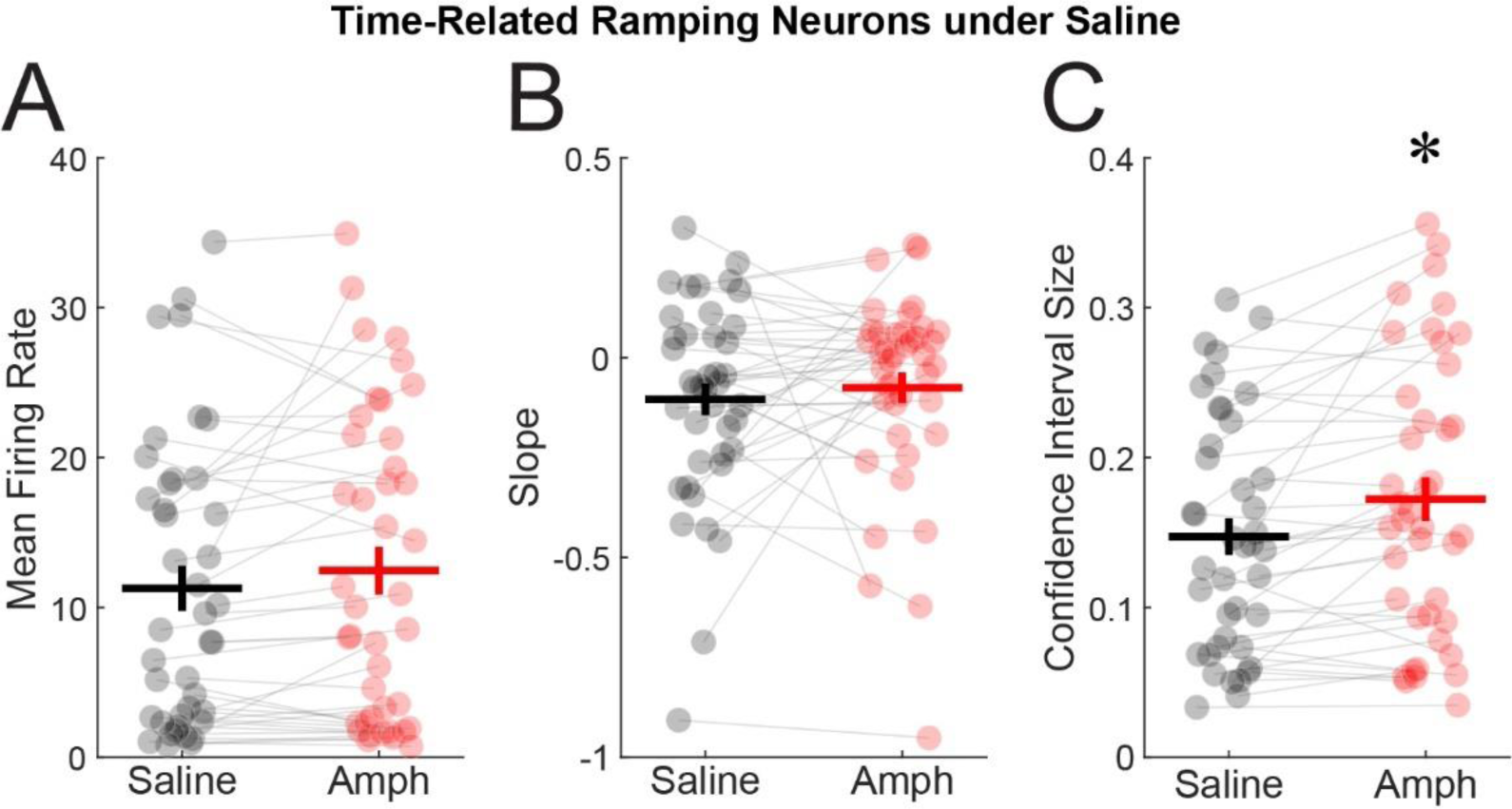
Amphetamine increases trial-by-trial variance in time-related ramping neurons. Amphetamine exposure did not change **(A)** mean firing rate or **(B)** estimated trial-by-trial slope from generalized linear models but **(C)** did increase trial-by-trial confidence interval size. Each dot represents a single neuron. The horizontal line represents the mean, and the vertical line represents standard error. * *p* < 0.05. Data from 41 neurons in 7 mice.

To ensure that this pattern held across our entire ensemble, we examined all 90 prefrontal neurons and found no difference in mean firing rate (saline: 10.63 ± 1.03 vs. amphetamine: 11.38 ± 1.17; F(_1,89_) = 2.03, *p* = 0.158; Fig. S2B) and no difference in trial-by-trial slope (saline: -0.05 ± 0.02 vs. amphetamine: -0.02 ± 0.02; F(_1,89_) = 2.12, *p* = 0.149; Fig. S2C), but we did find increased trial-by-trial CI size (saline: 0.14 ± 0.01 vs. amphetamine: 0.16 ± 0.01; F(_1,89_) = 13.83, *p* = 0.0004; Cohen’s *d* = 0.201; Fig. S2D; CI size increased in 63 of the 90 neurons, with 48 neurons increasing by more than 10%). Taken together, these data suggest that amphetamine increases the trial-by-trial variability of time-dependent ramping without changing the firing rate or ramp slope of prefrontal neurons. Because ramping activity is a mechanism of temporal encoding by prefrontal ensembles, these data provide insight into how amphetamine might impair interval timing behavior.

### Amphetamine disrupts functional interactions between time-related neurons

Joint peristimulus time histograms (JPSTHs) are useful to examine functional pairwise interactions between neurons (15,40). Under the saline condition, we found dynamic changes in functional correlations between pairs of ramping neurons over the interval during switch trials (Fig. 6A). Importantly, we found that amphetamine administration altered this relationship (Fig. 6B & C). Across 140 simultaneously-recorded, unique pairs of ramping neurons, we found JPSTH interactions were significantly decreased following amphetamine administration during the first 4 seconds of the interval (saline: 0.33 ± 0.05 vs. amphetamine: 0.10 ± 0.05; F(_1,139_) = 10.89, *p* = 0.001; Cohen’s *d* = 0.401; Fig. 6D). However, amphetamine did not change JPSTH interactions among the 163 simultaneously-recorded unique pairs of non-ramping neurons during the same interval (saline: -0.07 ± 0.03 vs. amphetamine: - 0.11 ± 0.03; F(_1,162_) = 0.88, *p* = 0.351; Fig. 6C). These data show that in addition to increasing trial-by-trial ramping variability, amphetamine decreased functional connectivity among prefrontal ramping neurons.

**Figure 6.**
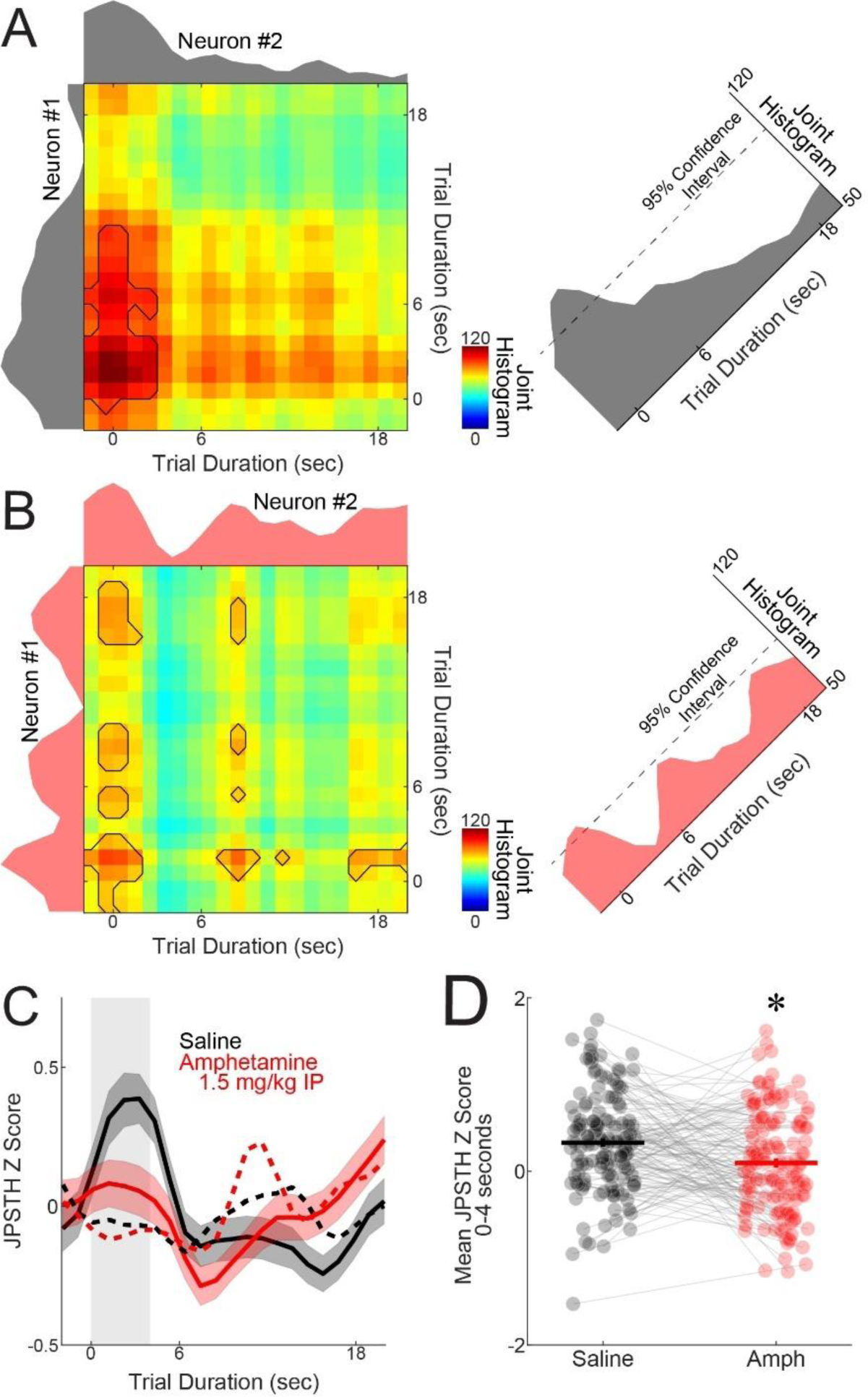
Amphetamine attenuates interactions between pairs of time-related ramping neurons. To examine pairs of time-related ramping neurons, we constructed joint peristimulus time histograms (JPSTHs) of trial-by-trial and bin-by-bin interactions. **(A)** A pair of neurons that displays time-related ramping activity under the saline condition. **(B)** The same pair of neurons under the amphetamine condition. **(C)** JPSTH interactions during the interval for ramping neurons under saline (black; data from 140 pairs of 41 ramping neurons) and matched pairs under amphetamine (red). Shaded area is standard error. Dashed lines are non-ramping pairs under saline and amphetamine (data from 163 pairs of 49 non-ramping neurons). **(D)** Mean JPSTH z-score from 0–4 seconds after trial start. Each dot represents a single pair of ramping neurons. The horizontal line represents the mean, and the vertical line represents standard error. * *p* < 0.05

### Amphetamine attenuates low-frequency cortical rhythms

Our prior work has highlighted the importance of frontal cortical low-frequency oscillations, ranging between 1–8 Hz, in both rodents and humans during interval timing; importantly, these low-frequency oscillations are dampened in rodents with disrupted dopamine and in patients with neurological and psychiatric disease (13,14,16,42,55,56). Consistent with these prior results, we found attenuated low-frequency 2–5-Hz power in rodents following amphetamine administration (saline: 1.92 ± 0.37 vs. amphetamine: -0.40 ± 0.66; F(_1,6_) = 6.32, *p* = 0.046; Cohen’s *d* = 1.685; Fig. 7A–D).

**Figure 7.**
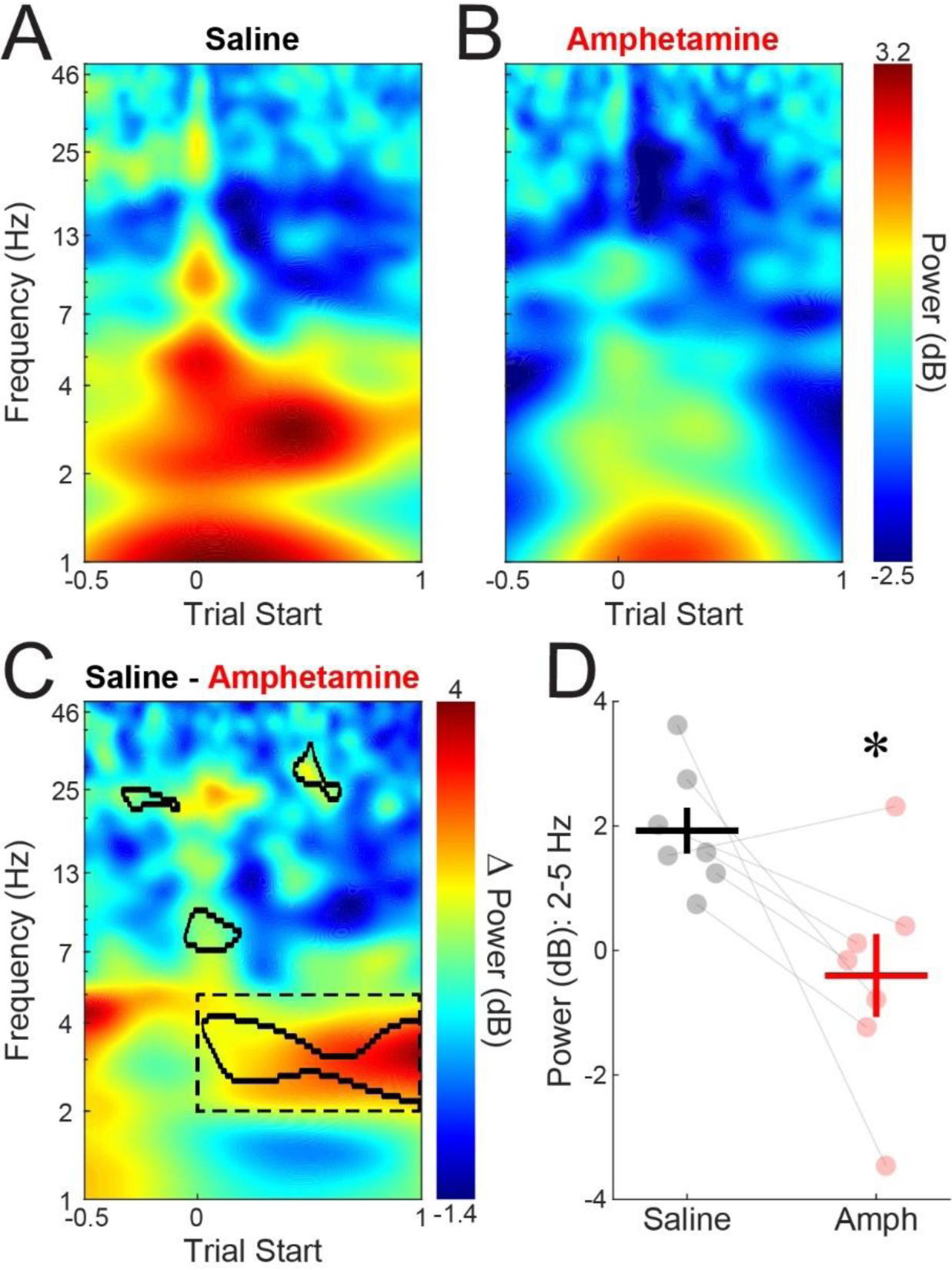
Amphetamine dampens low-frequency cortical rhythms. **(A)** Time-frequency analysis showed a burst of low-frequency 2–5-Hz power at trial start under the saline condition. **(B)** This low-frequency burst is decreased following amphetamine exposure. **(C)** Areas outlined with solid black lines indicate cluster sizes of at least 500 pixels with *p* < 0.05 via t-test, and the dashed lines indicate the time-frequency region of interest (2–5 Hz, 0–1 second). Subtraction of the saline minus amphetamine data revealed greater low-frequency power after trial start under the saline condition. **(D)** Power for the time-frequency region of interest showed a significant decrease in low-frequency power after amphetamine administration. Each dot represents a single mouse. The horizontal line represents the mean, and the vertical line represents standard error. * *p* < 0.05

## Discussion

We tested the hypothesis that amphetamine disrupts interval timing by degrading temporal encoding by prefrontal neurons. A meta-analysis of prior work revealed that acute amphetamine administration alters both timing precision and accuracy of interval timing behavior in rodents, with larger effects on timing precision. We found similar effects with acute administration of 1.5 mg/kg *_D-_*amphetamine during a mouse-optimized interval timing switch task. During this interval timing task, we recorded prefrontal activity and report three main results: acute amphetamine 1) increased the variability of prefrontal ramping; 2) attenuated functional interactions between pairs of ramping neurons; and 3) dampened low-frequency 2–5-Hz delta/theta rhythms in the prefrontal cortex. These data provide insight into how amphetamine disrupts prefrontal neuronal ensembles to impair interval timing behavior, predominately by increasing timing variability.

Amphetamine acts on catecholaminergic neurons in brain regions that are critical for interval timing, such as the prefrontal cortex. Here, we report evidence that acute amphetamine degrades time-related ramping and functional interactions among ramping neurons in the rodent prefrontal cortex. Our GLM analysis found more trial-by-trial variability in ramping slope, which resulted in fewer functional interactions among pairs of ramping neurons, as quantified by JPSTH (57). Because there is strong functional connectivity between ramping neurons early in the interval (15), increased trial-by-trial variability and decreased functional connectivity might contribute to interval timing impairments. These data provide insight into the mechanism of how amphetamine affects neuronal networks during interval timing.

Prefrontal neurons can encode temporal information through time-dependent ramping activity (13,22,39,58). This pattern lends itself to drift-diffusion computational models, in which neuronal spikes represent temporal evidence drifting toward a “threshold” (59,60). According to this model, increasing trial-by-trial variability in ramping slope would result in drift-diffusion computations crossing threshold at more variable times on a trial-by-trial basis, which would decrease timing precision. Here, we found evidence for 1) increased variability in ramping slope and 2) increased timing variability following acute amphetamine administration. These data provide one explanation for how amphetamine impairs interval timing. While there are other models of neuronal correlates of interval timing based on frequency dynamics (61) and distributed coding (62), our data suggest that changes in prefrontal ramping dynamics underlie the effect of amphetamine on higher-order executive functions.

Of note, amphetamine can also change dopaminergic signaling in another brain region: the striatum (63,64). Whereas striatal dopaminergic manipulations reliably affect timing accuracy (65,66), we find that amphetamine degrades both timing accuracy and precision, with larger effects on timing precision. Past work by our group and others have found that temporal encoding in the striatum is controlled by prefrontal input (15,67). It is possible that amphetamine also degrades temporal processing in the striatum, and our work suggests that these effects may be responsible for changes in timing accuracy. Future studies will explore this issue by combining acute amphetamine administration with striatal neurophysiology.

Amphetamine can impair executive functions in humans (4–8). These deficits in executive functions may contribute to poor decision-making, which in turn reinforces substance abuse behaviors and leads to relapse (68). Amphetamine has been reported to improve some executive functions (69,70), although these effects have been small and unreliable (71) and may be more relevant to disease populations, such as attention-deficit/hyperactivity disorder (ADHD) and schizophrenia (1,69). Amphetamine can also affect other nonspecific behavioral factors such as impulsivity, attention, and motivation (70,72–74). Catecholamines, especially dopamine, can have inverted-U-shaped nonlinear dynamics related to executive functions (28,75). Despite these complexities, our meta-analysis finds reliable evidence that amphetamine increases interval timing variability, which we also observed in our present study.

In addition to changes in ramping activity, we found that amphetamine changes prefrontal delta/theta oscillations between 2–5 Hz. These oscillations are an established mechanism of cognitive control (76), and our past work highlights how these oscillations can specifically engage ramping neurons via D1-dopamine receptors (D1DRs) (13,22,54). Amphetamine can trigger marked increases in prefrontal dopamine (77,78), which may alter signaling at prefrontal D1DRs and impair ramping activity. These data have translational relevance, as these oscillations are conserved between rodents and humans, can be detected via scalp electroencephalography (13,56,79), and can alter reward processing (80). In addition, delta/theta rhythms are specifically altered in human diseases such as Parkinson’s disease (13,14,55), schizophrenia (56,81), and ADHD (82). These data could lead to the development of neurophysiological markers of the effect of amphetamine on the prefrontal cortex.

Our findings have several limitations. 1) First, peripheral amphetamine administration is non-specific and will increase dopamine across the brain, including other brain regions involved in temporal processing, such as the striatum (65). Amphetamine also increases norepinephrine levels in several brain regions, including the prefrontal cortex (78,83,84). Future studies will need to study the effects of local amphetamine infusion in the prefrontal cortex and other relevant brain regions, such as the striatum, along with its effects on specific catecholamine receptor subtypes. 2) Second, we only used one 1.5 mg/kg dose of amphetamine, which was within the higher range of previous reports (Fig. 1). 3) Third, we chose to use forced drug administration, as in previous work, rather than a self-administration protocol to better understand how amphetamine interacts with prefrontal cortex neuronal ensembles. 4) Finally, because our recordings sampled only a small number of neurons and a small area of the prefrontal cortex, it is challenging to relate our measures of neuronal activity from a small section of one prefrontal cortical hemisphere to overall interval timing behavior. Future work will need to study other doses and other drug administration protocols to better understand the effects of amphetamine on cognitive processing.

In summary, we demonstrated that amphetamine decreases temporal precision by degrading prefrontal temporal encoding. Our meta-analysis shows that amphetamine reliably degrades temporal precision across prior studies, and we find that acute amphetamine increases trial-by-trial variability of time-related ramping, decreases functional interactions among pairs of prefrontal ramping neurons, and attenuates prefrontal delta/theta activity. These data provide novel insight into how amphetamine affects prefrontal networks, which may be relevant for understanding amphetamine as a drug of abuse and developing new therapies that target the prefrontal cortex and treat amphetamine use.

## Authorship

MAW, KS, HRS, and NSN designed the experiments. MAW, KS, BQK, and EET performed all experiments. MAW, ASB, YK, and NSN performed all statistical analysis. MAW, KS, EET, and NSN wrote the manuscript, and all authors reviewed and revised the manuscript. We gratefully acknowledge Heather Widmayer, MS, MBA of the Scientific Editing and Research Communication Core at the University of Iowa Carver College of Medicine for suggestions toward improving the clarity of the manuscript.

## Data and code availability

All data and code are available at https://narayanan.lab.uiowa.edu/article/datasets.

## Supplemental Information

**Supplemental Figure 1.**
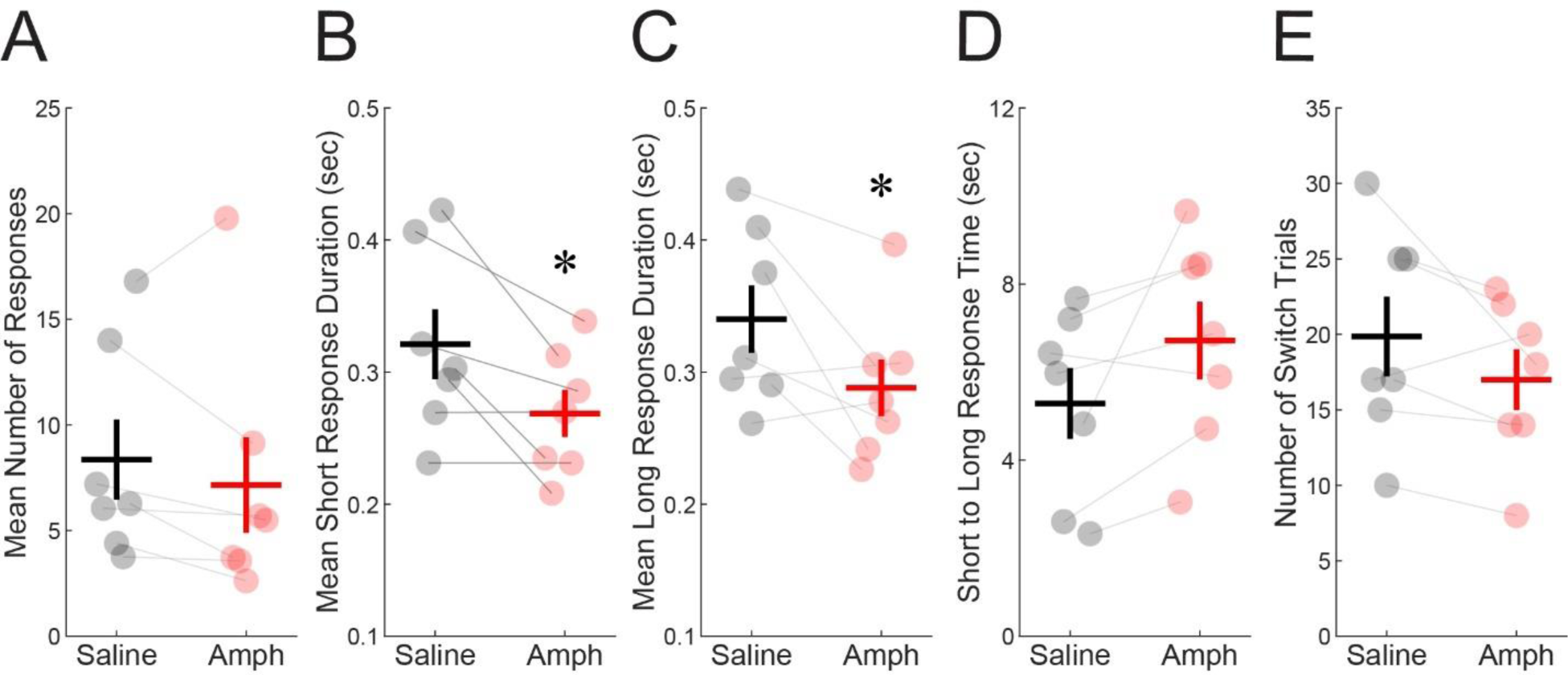
Amphetamine does not change the mean number of responses per trial **(A)** but does decrease the mean response duration at the short **(B)** and long **(C)** nosepoke ports. Amphetamine does not change the time between the final short and first long responses during switch trials **(D)** or the number of switch trials **(E)**. Each dot represents a single mouse. The horizontal line represents the mean, and the vertical line represents standard error. * *p* < 0.05

**Supplemental Figure 2.**
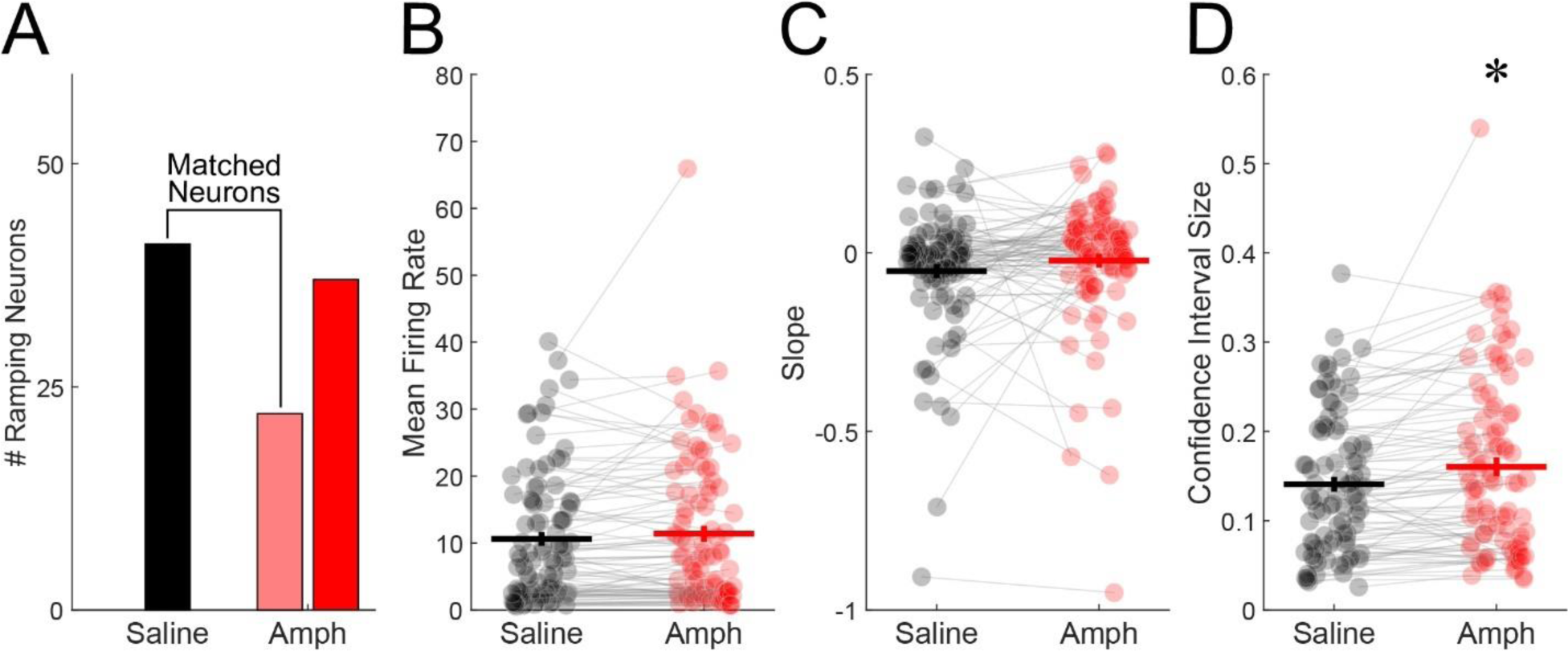
Amphetamine affects all prefrontal neurons. **(A)** Out of 90 prefrontal neurons, 41 (∼46%) displayed time-related ramping activity under saline (black). Following amphetamine administration, only 22 of those 41 neurons still displayed time-related ramping activity (light red). However, a total of 37 neurons exhibited time-related ramping activity under amphetamine (red), meaning that 15 of those neurons did not exhibit time-related activity under saline. **(B–D)** Analysis of all prefrontal cortex neurons. Amphetamine did not change mean firing rate **(B)** or estimated trial-by-trial slope **(C)** from generalized linear models but did **(D)** reliably increase trial-by-trial confidence interval size. Each dot represents a single neuron. The horizontal line represents the mean, and the vertical line represents standard error. * *p* < 0.05

